# Molecular dynamics simulations of the Spike trimeric ectodomain of the SARS-CoV-2 Omicron variant: structural relationships with infectivity, evasion to immune system and transmissibility

**DOI:** 10.1101/2022.02.14.480347

**Authors:** Vitor Martins de Freitas Amorim, Robson Francisco de Souza, Anacleto Silva de Souza, Cristiane Rodrigues Guzzo

## Abstract

The severe acute respiratory syndrome coronavirus 2 (SARS-CoV-2) Omicron variant is replacing Delta, the most prevalent variant worldwide from the beginning of 2021 until early 2022. The Omicron variant is highly transmissible and responsible for a new worldwide COVID-19 wave. Herein, we calculated molecular dynamics simulations of the SARS-CoV-2 trimeric spike protein of Wuhan-Hu-1 strain (wild type, WT) and the Omicron variant of concern. Structural analyses reveal that the Spike^Omicron^ presents more conformational flexibility than Spike^WT^, mainly in the N-terminal domain (NTD) and receptor-binding domain (RBD). Such flexibility results in a broader spectrum of different conformations for Spike^Omicron^, whereby the RBD can more easily visit an up-conformational state. We reported how the mutations in this variant may influence the intra- and inter-protomer contacts caused by conformational flexibility of the NTD. Based on our analysis, we suggest that the differences in conformational flexibility between Spike^Omicron^ and Spike^WT^ may explain the observed gains in infectivity, immune system evasion and transmissibility in this novel variant.

**Graphical abstract:** 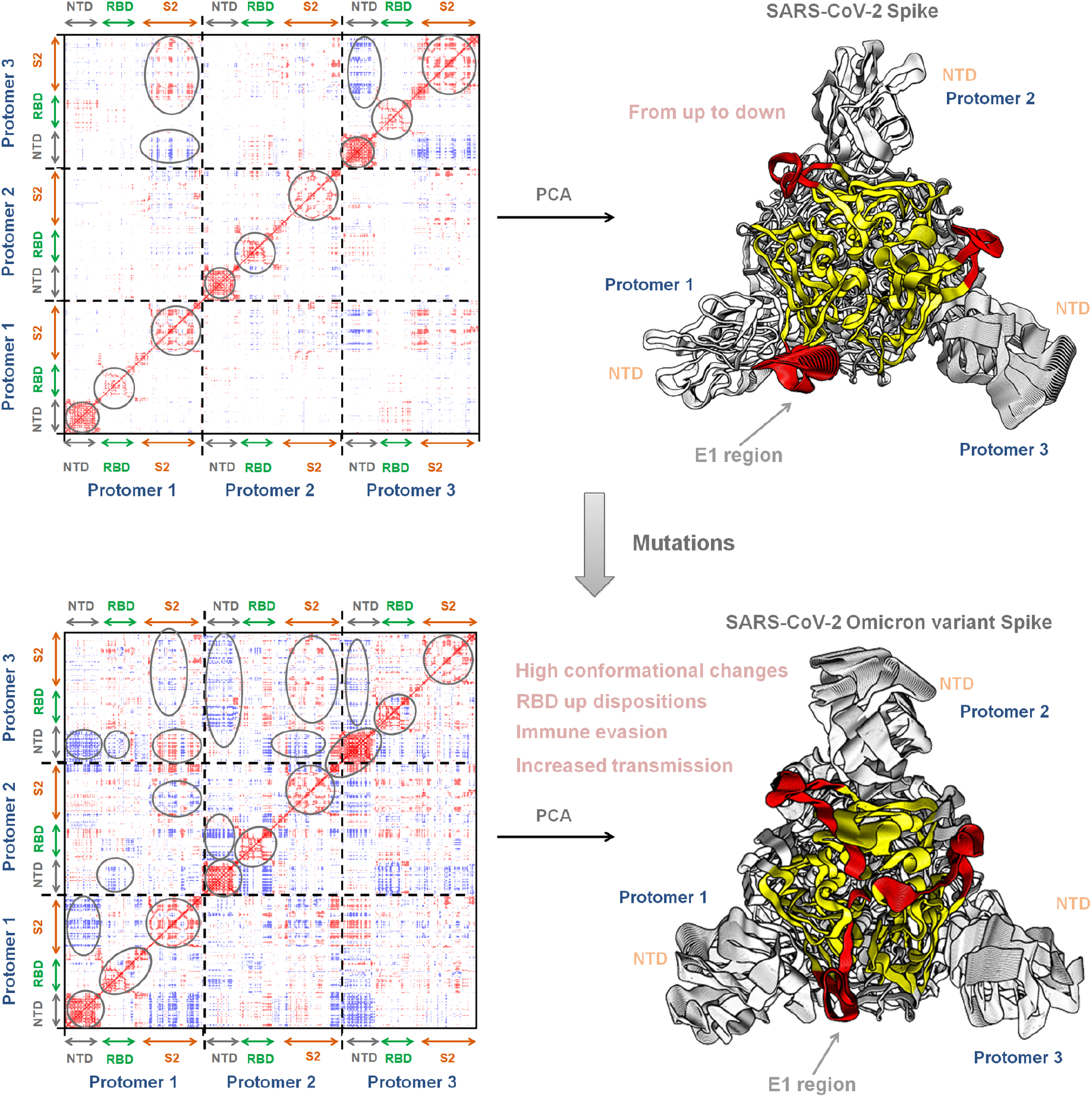

## Introduction

Despite efforts in vaccination in several countries, more than 400 million cases and 5.8 million deaths by Severe Acute Respiratory Syndrome Coronavirus 2 (SARS-CoV-2) have been reported since the beginning of the Coronavirus diseases 2019 (COVID-19) pandemic [1]. Since the start of the pandemic, several SARS-CoV-2 variants of concern (VOC) as the Alpha (B.1.1.7), Beta (B.1.351), Gamma (P.1), Delta (B.1.617.2) and Omicron (B.1.1.529) emerged and propagated worldwide, all showing higher transmission rates than the original Wuhan-Hu-1 strain (wild type, WT). In general, the viral genome encodes four structural proteins such as Spike protein (or S protein), nucleocapsid (N), membrane (M) and envelope (E). Furthermore, it contains 16 non-structural proteins [2] (**Figure 1a**). In particular, the trimeric Spike ectodomain has 1,273 amino acids containing two subunits, S1 and S2 (**Figure 1b-c**). The S1 subunit contains the N-terminal domain (NTD), a receptor-binding domain (RBD) and SD1 and SD2 domains. The S2 subunit is composed by the fusion peptide (FP), the heptapeptide repeat sequences 1 (HR1) and 2 (HR2), a transmembrane domain (TM) and C-terminal domain (CT) [3,4]. The functional role of S1 is related with the NTD and RBD, which the first acts as a “wedge”, playing a key role in the structural changes of the trimeric Spike protein [5] while RBD interacts with human angiotensin-converting enzyme 2 (hACE2) (**Figure 1d**). The S2 subunit favors fusion between viral envelope and host cell membrane, which is mediated by domains HR1, HR2, TM and CT.

**Figure 1.**
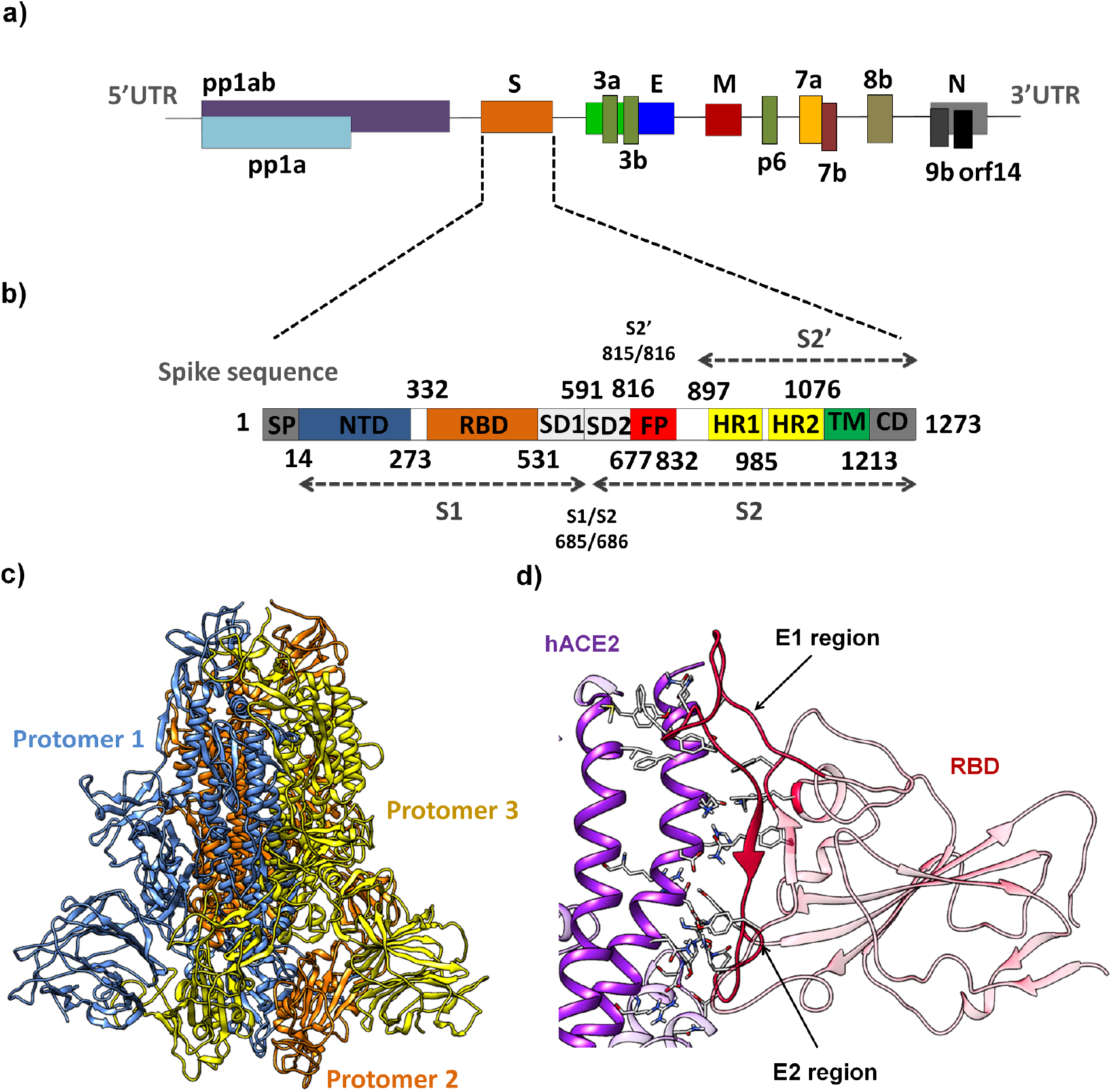
representative schematic of the amino acid sequence and the three-dimensional structure of the Spike protein. **a)** Polyprotein of SARS-CoV-2. **b)** Spike sequence showing the S1 and S2 subunits, NTD, RBD, SD1, SD2, FP, HR1, HR2, TM and CD. c) Tridimensional structure of the trimeric Spike ectodomain (PDB ID 7CAB[10]). d) RBD interacting with hACE2 (PDB ID 6M0J[11]). The painel d) describes two regions, E1 and E2. E1 region is delimited by K417, L455, F456 and residues 470-490[12]. The E2 region corresponds to residues 446-453 and 493-505 residues[12].

The Alpha variant has several mutations in the Spike protein such as Δ69-70, Δ144, N501Y, A570D, D614G, P681H, T716I, S982A and D1118H. The Beta variant has mutations L18F, D80A, D215G, Δ242-244, R246I, K417N, N501Y, D614G and A701V [6]. The Gamma variant has mutations L18F, T20N, P26S, D138Y, R190S, K417T, E484K, N501Y, D614G, H655Y, T1027I and V1176F [6]. The Delta variant has mutations T19R, Δ157-158, L452R, T478K, D614G, P681R and D950N [6]. The L452R substitution in the Spike protein can make the novel coronavirus 50% more transmissible than the Alpha variant, in addition to conferring better infectivity due to favorable electrostatic interactions with hACE2 [7]. Although the position of the T478K substitution is far from the hACE2-interacting interface, K478 mediates better interaction with hACE2 than T478 by a compensation mechanism to form stable conformations of the Spike/hACE2 complex [8,9]. Omicron variant has many mutations in Spike protein such as A67V, Δ69-70, T95I, Δ142-144, Y145D, Δ211, L212I, ins214EPE, G339D, S371L, S373P, S375F, K417N, N440K, G446S, S477N, T474K, E478K Q493R, G496S, Q498R, N501Y, Y505H, T547K, D614G, H655Y, N679K, P681H, N764K, D796Y, N856K, Q954H, N969K and L981F. This variant spreads further than the ancestral SARS-CoV-2, becoming predominant over the Delta variant worldwide [6].

In particular, the Omicron variant Spike has insertions (EPE at position 214) and deletions (Δ69-70, Δ142-144, Δ211) [13,14]. In SARS-CoV-2 VOCs, the mutations in the Spike protein often cause an increase in the binding affinity to hACE2 [15,16]. However, when these binding affinity values are compared between VOCs, they are within the same order of magnitude [17–20], suggesting that transmissibility, viral load and lethality may also be related to other molecular factors, such as: (1) an increase of RBD up dispositions in Spike [18,19,21]; (2) immune system evasion [22–26]; and (3) level of exposure of the furin cleavage site to proteases [27–29].

The loss of sensibility to neutralizing antibodies against Omicron may be explained due to the Δ144-145 in the NTD [20]. We hypothesize that insertions and deletions observed in this domain could also be associated with further structural changes between the protomers within the trimeric Spike protein. This hypothesis is supported by the observation that structural changes in the adjacent loops 246—260 and nearby loops 144—155 may decrease recognition of neutralizing antibodies [28]. In addition, P681H mutations near the furin cleavage site (residues 682-685) may slightly increase S1/S2 cleavage [29], probably because this substitution enhances exposure of this site to proteases as it confers greater structural flexibility to this region and probably decrease thermostability of proteins. Proline is the most rigid residue and substitution of this residue in the protein may cause a gain in local flexibility [30]. Furthermore, the D614G mutation may also increase the cleavage by furin, by changing the Spike ectodomain structure caused by an allosteric effect [31].

In order to investigate how the mutations found in the Omicron variant affect its trimeric structure in relation to the Spike of Wuhan-Hu-1 strain (sequences in **Figures S1** and **S2**), we performed molecular dynamics simulations of the trimeric ectodomains of the Spike^WT^ and Spike^Omicron^ for 100 ns. We used root-mean-square deviation (RMSD), root-mean-square fluctuation (RMSF), principal component analysis (PCA) and hydrogen bonding occupancies for investigating the structural aspects associated with the mutations of this variant. We observed that the Spike^Omicron^ presents more conformational flexibility than Spike^WT^, mainly in the NTD and RBD. A broader spectrum of different conformations for Spike^Omicron^ may be influenced by the intra- and inter-protomer contacts caused by conformational flexibility of the NTD. Based on our analysis, we suggest that the differences in conformational flexibility between Spike^Omicron^ and Spike^WT^ may explain the observed gains in infectivity, immune system evasion and transmissibility of the Omicron variant.

## Materials and Methods

### Homology modeling

To predict the SARS-CoV-2 Spike structure of WT and Omicron variant, we performed homology modeling using the primary sequences of amino acids corresponding to WT (PDB 7CAB_A[10]) and B.1.1.529 (Omicron variant, accession number UFO69279.1). All sequences were submitted in the Swiss-Model web-service[32–34], ignoring the transmembrane regions. The final sequences are shown in **Figure S1.** Using the <u>B</u>asic <u>L</u>ocal <u>A</u>lignment <u>S</u>earch <u>T</u>ool (BLAST)[35], we considered the Spike^Omicron^ mutations indicated in **Figure S2**. From the generated structures, we selected the down conformation in both proteins as input for molecular dynamics simulations (models 1 and 2, corresponding to Spike^WT^ and Spike^Omicron^, respectively, are available in **supplementary file**).

### Setup for molecular dynamics simulations

We performed molecular dynamics simulations to study the structural features of the Spike trimer, considering all 3D structures modeled in Swiss-Model. We determined the protonation states of ionizable residues in an implicitly aqueous environment at pH 7.0 using the PROPKA module implemented in the Maestro program (academic version v. 2021-2, by Schrödinger)[36]. Thus, all glutamic and aspartic residues were represented as non-protonated, arginine and lysine residues were assumed to have a positive charge, in the N- and C-terminal regions, the amino and carboxyl groups have been converted to charged groups. The histidines were protonated according to prediction of PROPKA and manual inspection, as: **WT**, δ-tatomer (H66, H146, H519, H655, H1058, H1064, H1088) and ε-tatomer (H49, H69, H207, H245, H625, H1048, H1083, H1101); **Omicron**, δ-tatomer (H66, H141, H202, H502, H516, H678, H1085) and ε-tatomer (H49, H242, H622, H1045, H1061, H1080, H1098) and charged histidines (H951 and H1055).

All molecular dynamics simulations were performed in GROMACS, v. 5.1.5[37–40], using the OPLS-AA force field[41]. All systems were then explicitly solvated with TIP3P water models in a cubic box and neutralized, maintaining the concentration of 150 mM NaCl and minimized until reaching a maximum force of 10.0 kJ/mol or a maximum number of steps in 50,000. The systems were consecutively equilibrated in isothermal-isochoric (NVT) ensembles by 5 ns (number of steps and intervals of 2,500,000 and 2 fs, respectively), isobaric ensemble using annealing (0 to 320 K, 0.1 ps steps) by 5 ns (number of steps and intervals of 2,500,000 and 2 fs, respectively) and isothermal-isobaric (1 bar, 310 K, NpT) by 5 ns (number of steps and intervals of 2,500,000 and 2 fs, respectively).

All simulations were then performed in a periodic cubic box considering the minimum distance of 1.2 nm between any protein atom and the walls of the box. Molecular dynamics runs were performed for 100 ns (number of steps and intervals of 50,000,000 and 2 fs, respectively) in the NpT ensemble. We use the leap-frog algorithm to integrate Newton’s equations. We selected LINCS (LINEar Constraint Solver) that satisfies the holonomic constraints, whose number of iterations and order were 1 and 4, respectively. We use neighbor search grid cells (Verlet clipping scheme, frequency to update 20-step neighbor list, and clipping distance for 12 Å^2^ short-range neighbor list). In the van der Waals parameters, we smoothly shift the forces to zero between 10 and 12 Å. In electrostatic Coulomb, we use Particle-Mesh Ewald (SPME) fast and smooth electrostatics for long-range electrostatics. In addition, we set the distance for the Coulomb cut to 12 Å, order of interpolation for PME to value 4 (cubic interpolation) and grid spacing for Fast Fourier Transform (FFT) to 1.6 Å. In temperature coupling, we use velocity rescheduling with a stochastic term (V-resale, modified Berendsen thermostat). After obtaining two coupling groups (protein and water/ions), we consider the time constant (0.1 ps) and 310 K as the reference temperature. In the pressure coupling (NpT ensembles), we use the Parrinello-Rahman barostat (isotropic type, time constant of 2 ps, reference pressure of 1 bar and isothermal compressibility of 4.5×10^-5^ bar^-1^). In molecular dynamics calculations, we use periodic boundary conditions in xyz coordinates (3D space). We then calculated the percent hydrogen bond occupancy (all frames, considering cut-off of distance and angle of 4 Å and 20 degrees, respectively), root-mean-square deviation (RMSD) and root-mean-square fluctuation (RMSF) using the GROMACS modules and virtual molecular dynamics (VMD)[42]. Analysis of C_α_ cross-correlated displacements were performed using the R-based package Bio3d[43].

### Principal component analysis

Principal component analysis was calculated using the GROMACS package. Using simulated molecular dynamics trajectories, we determine the average position of the Spike backbone’s atoms and calculate correlation matrix fitting non-mass weighted. Correlation matrices were constructed using the Spike^WT^ (30,537×30,537) and Spike^Omicron^ (30,375×30,375) trimer backbone. In both, only 10,000 eigenvalues were saved.. Using first and second eigenvalues, we determined the eigenvectors 1 and 2 (named PC1 and PC2). PC1 represents the first main movement detected along molecular dynamics trajectories. PC2 represents the second main movement detected in molecular dynamics. From PC1 and PC2, we observed 300 snapshots corresponding to MD trajectories projected in these eigenvectors (**supplementary movies S1**-**S8**). We used first and second eigenvectors to verify trajectory scores of the Spike^WT^ and Spike^Omicron^ and compare with the respective conformational states.

## Results and Discussion

The root-mean-square deviation (RMSD) of the Spike^Omicron^ and Spike^WT^ protomers are shown in **Figure S3**. In general, Spike^Omicron^ presents more conformational changes) than Spike^WT^ (**Figures S1, S4** and **S5**). These structural changes may be related with high flexibility of residues localized mainly in the S1 subunit (NTD and RBD) (**Figure 2**). We observed a decrease in flexibility in the furin cleavage region in the Omicron variant compared to the WT (**Figure 2**). However, the structural model of the Spike^WT^ for the MD analysis was based on the structure with PDB ID 7CAB that has a “GSAS’’ instead of “RRAR” at the furin cleavage site [10]. The root-mean-square fluctuation (RMSF) of Spike^WT^ presented high fluctuations in the NTD (residues 69-77, 144-155 and, 246-258), in the RBD (residues 470-490) and furin cleavage region (residues 676-690). The Spike^Omicron^ showed flexibility in the NTD (residues 70-258), in the RBD (residues 466-498), in SD2 (residues 615-638) and, in the furin cleavage region (residues 673-687). In the Spike^Omicron^, the NTD mutations, deletions and insertions (A67V, Δ69-70, T95I, Δ142-144, Y145D, Δ211, L212I, insertion of EPE at position 214) may favor the up-conformation of the RBD as well as in the increasement of flexibility for this domain in relation to SARS-CoV-2^WT^. Indeed, the structure of Spike^Omicron^ by cryo-EM revealed that the NTD region is more flexible when compared with the remaining protein [20].

**Figure 2.**
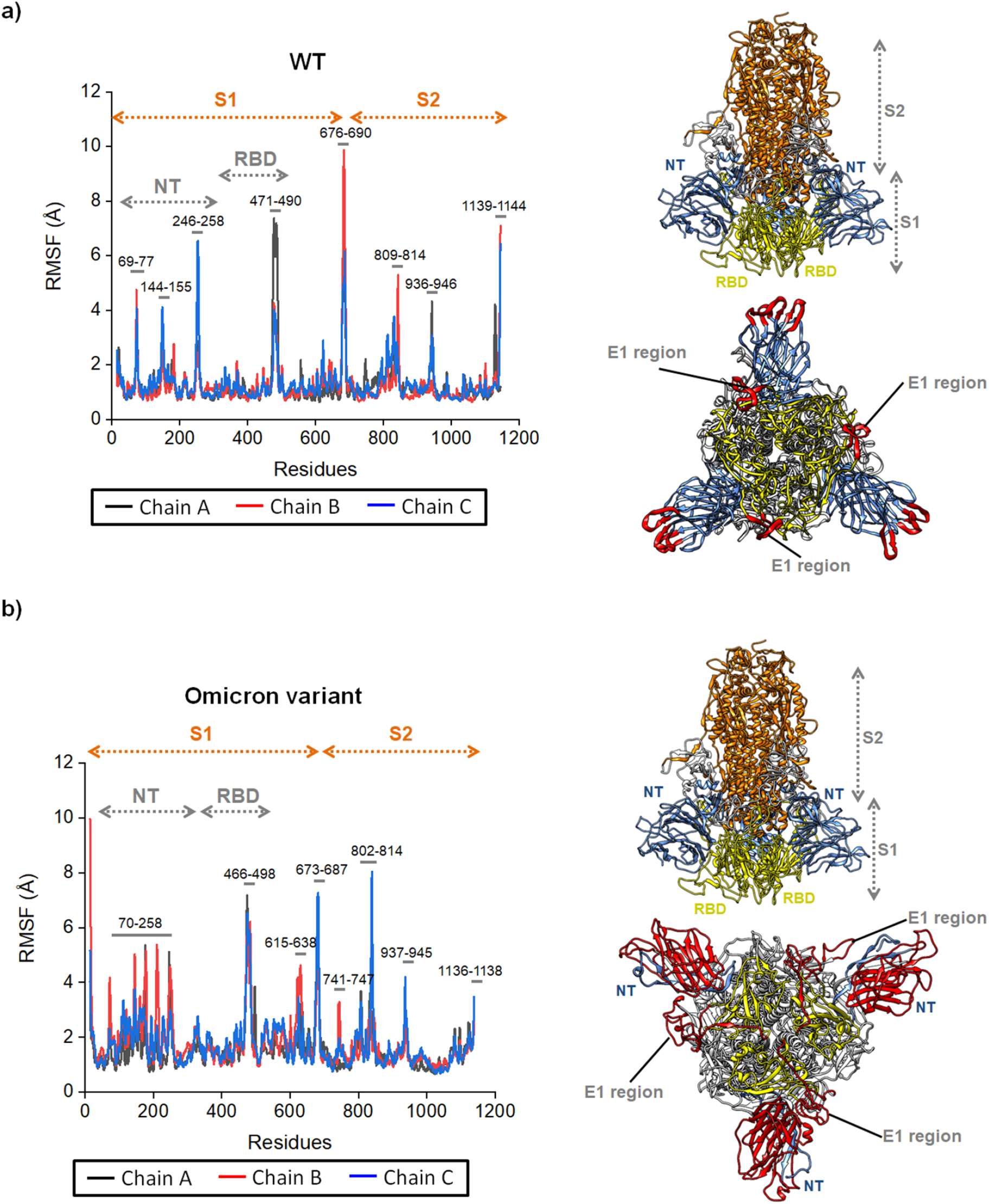
Structural aspects obtained by molecular dynamics simulations of Spike^WT^ and Spike^Omicron^. Backbone root-mean-square fluctuation (RMSF) of **a)** Spike^WT^ and **b)** Spike^Omicron^. The Spike^WT^ presented high flexibility in NTD (residues 69-77, 144-155, 246-258), in RBD (residues 470-490) and S1/S2 region (residues 676-690). The Spike^Omicron^ has more flexibility in NTD (residues 70-258), in RBD (residues 466-498), in SD2 (residues 615-638) and S1/S2 interface (residues 673-687). Mutations (A67V, Δ69-70, T95I, Δ142-144, Y145D, Δ211, L212I, insertion of EPE at position 214) in the NTD contribute to RBD to adopt an up conformation in Spike^Omicron^.

**Figure 3.**
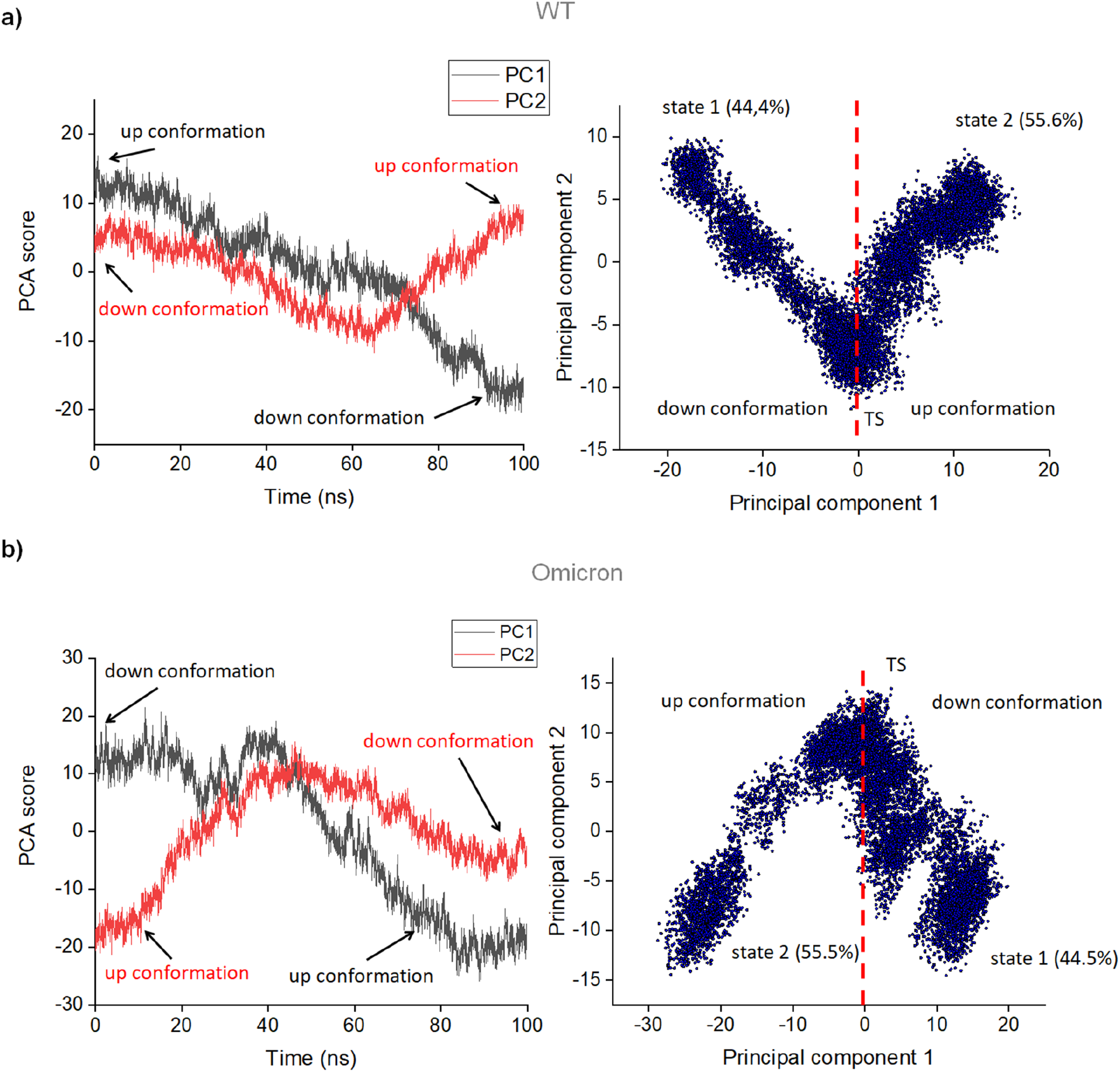
Principal component analysis (PCA) of the molecular dynamics trajectories. From the average structure of Spike backbone (WT or Omicron variant), the correlation matrix (fitting non-mass weighted) was calculated. After diagonalizing the correlation matrix, the eigenvalues were determined (**Table S1**) and used to extract the orthogonal eigenvectors (or principal components). The principal components represent the maximum correlation between residue pairs along the MD simulation. MD trajectories were projected in the eigenvector 1 (or PC1, principal component 1) to obtain the main movements of the Spike protein. Similarly, MD trajectories were projected in the eigenvector 2 (or PC2, principal component 2) to obtain the second main movements of the Spike protein. The PC2 movements are less significant than PC1 movements. From projected MD trajectories in the two first principal components, PCA scores were calculated. The PCA score as a function of time (left) and PCA score plot (right) were performed from the backbone of **a)** Spike^WT^ and **b)** Spike^Omicron^.

The RBD domain of the Spike protein binds human angiotensin-converting enzyme (hACE2) by two regions, E1 and E2 [12] (**Figure 1**). E1 binds hACE2 mainly through hydrophobic interactions. In contrast, the E2 region, made up of residues 444-454 and 493-505, mediates mostly polar interactions with hACE2 (**Figure 1d**). Curiously, we did not observe significant changes in RMSF in the residues affected by subtitutions G339D, S371L, S373P, S375F, K417N, N440K, G446S, N501Y, Y505H and T547K. However, the mutations S477N, T478K, E484A, Q493R, G496S, Q498R, localized in the RBD, correspond to the E1 region, which has intercalated hydrophobic and hydrophilic residues in WT [12]. The E1 region may be the first to be recognized by hACE2 and presents high RMSF values in the residues 470-490 [12].

Previous study has shown that the D614G mutation plays a key role in SARS-CoV-2’s evolution because it is involved with the disruption of the interprotomer contact [44], modulates structural conformations that promote membrane fusion [27], increases virion spike density and infectivity [45,46], and increases the cleavage rate by furin after causing conformational changes in the Spike ectodomain [31]. Accordingly, we also verified that the G614 residue influences the conformational space of the Spike^Omicron^ protein. Analysis of the trajectories of the Spike^WT^ residues shows that stable interactions between the SD2 subdomain of one protomer and the closest S2 subunit of a neighboring protomer are mediated by hydrogen bonding between residues Asp614—Met835 (hydrogen bonding occupancy of 41.8%), Asp614—Tyr837 (hydrogen bonding occupancy of 84.0%), and Asp614—Arg854 (hydrogen bonding occupancy of 99.8%) (**Figure S6**). Such interactions confer some degree of structural rigidity to the Spike^WT^ ectodomain. On the other hand, G614 does not interact with any amino acid residues in Spike^Omicron^ and the loss of the aforementioned interactions increases mobility of the SD2 domain and of the link between the FP and HR1, which contains the S2’ cleavage site, making the Spike^Omicron^ ectodomain more flexible. Although the G614 residue is distant from the RBD, this substitution may also affect the RBD up/down dispositions [31]. The Spike S1 subunit is involved in recognition of the host cell through receptor binding and the S2 subunit participates in membrane fusion [3,4]. Mutations near the S1/S2 boundary (residues 682-685) may be associated with increased efficiency of cleavage by furin in VOCs. Spikes of D614G, Alpha, Beta, and Delta variants are efficiently cleaved by furin between residues 682-685 [27–29]. In agreement with cryo-EM data for Spike ectodomain from different SARS-CoV-2 VOCs, our molecular dynamics data reveal that this region is highly flexible in Spike^Omicron^ (**Figures 1**).

To gain insights into the differences between the MD trajectories of Spike^WT^ and Spike^Omicron^, we performed a principal component analysis of the covariance matrix for positional deviations of backbone atoms of the Spike trimer [47]. From the diagonalization of the correlation matrices, we obtained eigenvalues and their corresponding orthogonal eigenvectors (**Table S1**). Projection of the original covariances onto the orthogonal space defined by the eigenvalues allows calculation of the eigenvectors. The highest eigenvectors are called principal components and represent the most important correlations between residues pairs. We evaluate the first and second eigenvectors, which represent the most relevant motions of the Spike protein detected by molecular dynamics trajectories. The PCA score plot reveals that the Spike protein visits two main sets of states throughout the simulation (**Figure 4**). Using the first eigenvector (PC1, first principal component), we observed that the most prominent movement of the Spike^WT^ is the well known repositioning of the RBD from the up- to the down-conformation (**Figure S7a** and **supplementary movies S1** and **S2**). The second eigenvector (PC2, second principal component), however, showed that the second most important movement of Spike^WT^ RBD is a reverse process, from down- to up-conformation state (**Figure S7b** and **supplementary movies S3** and **S4**). Therefore, PCA-projected trajectory suggests that Spike^WT^ RBD tends to visit more down than up conformation, exposing the residues 470-490 (E1 region) for interactions with hACE2.

**Figure 4.**
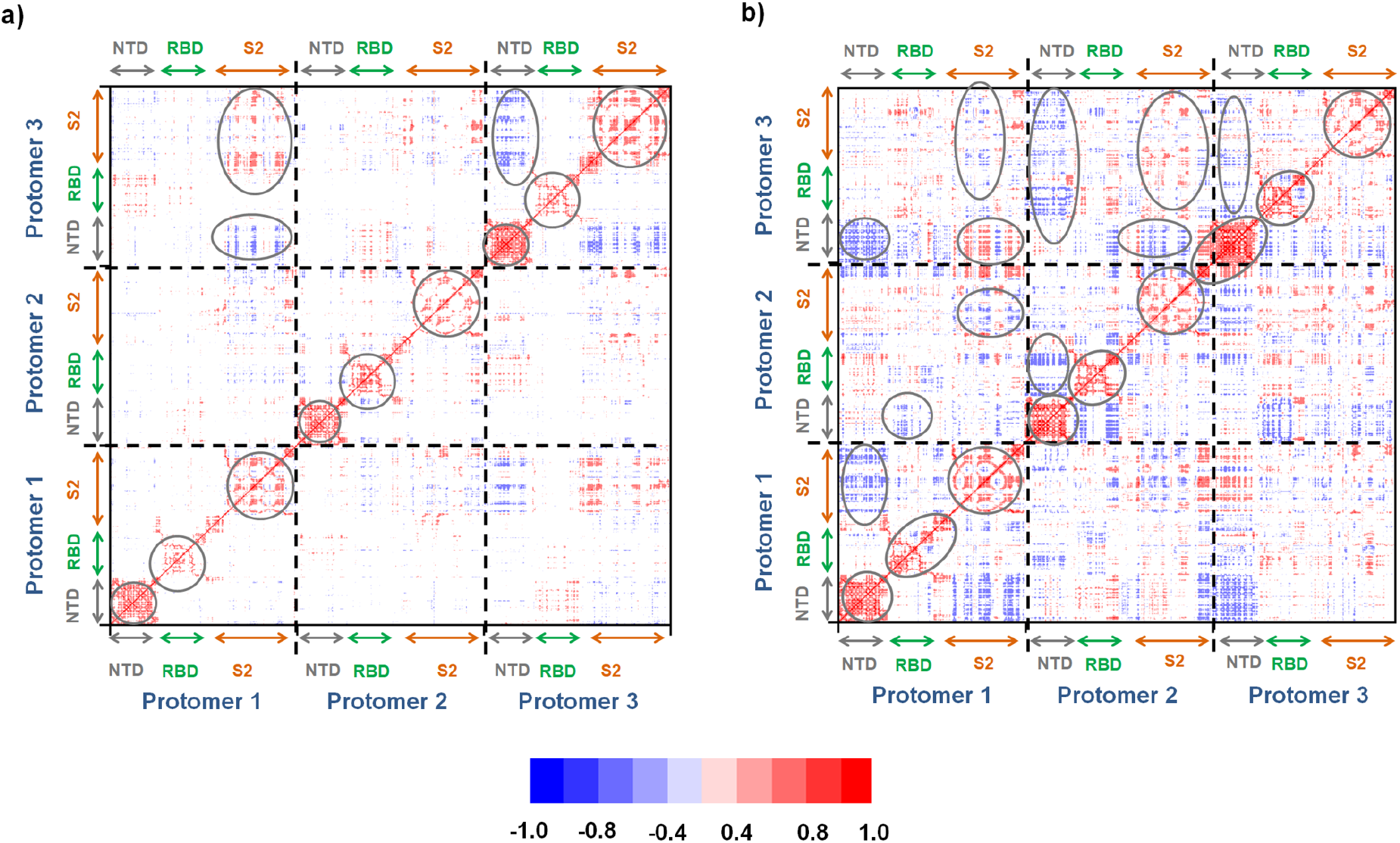
Trimeric Spike correlation matrix of the wild type and Omicron variant. The correlations were calculated to find correlated motions between internal residues and interprotomer residues in **a)** WT and **b)** Omicron. The gray circles indicate how the movement of residues influences the mobility of the Spike protein. The red and blue dots show the positively and negatively correlated pairs, respectively. Note that the correlations are more intenses in Spike^Omicron^ than Spike^WT^. In Spike^WT^, the negative and positive correlations are observed between pairs S2^protomer 1^/NTD^protomer 3^ and RBD^protomer 3^/S2^protomer^ 3, respectively. In Spike^WT^, all protomers were positively correlated between their domains (main diagonal matrix) but notably only protomer 3 presented anti-correlation between NTD/S2. In contrast, Spike^Omicron^ showed more anti-correlated residue pairs between protomers than Spike^WT^. The internal anti-correlations between their domains are more than WT. In Spike^Omicron^, the anti-correlation are observed between pairs NTD^protomer 1^/S2^protomer^, NTD^protomer 1^/NTD^protomer 3^, RBD^protomer 1^/RBD^protomer 2^, RBD^protomer 1^/RBD^protomer 3^, S2^protomer 1^/S2^protomer 2^, S2^protomer 1^/RBD^protomer 3^, S2^protomer 1^/S2^protomer 3^, NTD^protomer 2^/RBD^protomer 3^, NTD^protomer 2^/S2^protomer 3^ and S2^protomer 2^/NTD^protomer 3^. Conversely, only the pairs S2^protomer 1^/NTD^protomer 3^ and S2^protomer 2^/S2^protomer 3^ were positively correlated.

As expected, the PCA score plot also showed that Spike^Omicron^ RBD populates the up and down conformations but, according to PC1-projected trajectory, it visits preferably from down- to up-conformation state (**Figure S8a** and **supplementary movies S5** and **S6**). The reverse process to PC1-projected trajectory also was observed, with the Spike^Omicron^ RBD changing from the up to down conformation (**Figure S8b** and **supplementary movies S7** and **S8).** Interestingly, the NTD is more flexible in Spike^Omicron^ than Spike^WT^, probably because its domain suffered mutations and also insertions and deletions that probably caused significant structural rearrangement allowing it to increase the RBD up dispositions, as observed by previous studies [31]. In agreement with RMSF analysis (**Figure 1**) and hydrogen bonding occupancies, our PCA data suggest that the D614G is fundamental for improving the RBD up dispositions, contributing to increase the viral infection rate of the Omicron variant.

In order to investigate the effect of the mutations in the Spike^Omicron^, we also calculated the C_α_ correlation matrix of the trimeric Spike^WT^ and Spike^Omicron^. **Figure 4** shows the heat map of the correlation matrix, considering internal correlations and also the long-range interactions between each protomer in both proteins. In general, the anti-correlations may be associated with the movements observed in the PC1-projected trajectory, indicating that the RBD up dispositions on Spike^Omicron^ also may be governed by NTD movements. Mutations in NTD seem to play a functional role fundamental in the Spike, making the protein more flexible and efficient to infect the host cell than wild type. Furthermore, such greater flexibility probably favors structural rearrangements that difficulties neutralizing antibodies, contributing to the immune evasion. Indeed, the SARS-CoV-2 VOCs evade the neutralizing antibodies contained in plasma from convalescent or/and immunized patients [48] and both NTD and RBD carry important epitopes for recognition by neutralizing antibodies, the later one being the major target for these interactions [49]. Deletion of the residues L242, L243 and A244 in Spike^Beta^ causes structural changes in the adjacent loops 246—260 and nearby loops 144—155, decreasing recognition by neutralizing antibodies [28]. Since neutralizing antibodies show more sensitivity to recognize SARS-CoV-2^WT^ than SARS-CoV-2^Omicron^ and that the antibodies mainly recognize the NTD region and the RBD, we suggested that the significant conformational changes of the NTD and RBD of Spike^Omicron^ caused not only by deletions and insertions, but also by the D614G mutation, may be related to the loss of sensitivity of neutralizing antibodies. As shown in **Figure S8a** and **Figure 4**, the bigger structural movements in the NTD^Omicron^ and RBD^Omicron^ may be related with a steric hindrance during neutralization by antibodies, which may explain immune system escape. The loss of sensitivity of neutralizing antibodies for SARS-CoV-2 Omicron variant probably contributes to higher virus transmissibility than WT. Since D614G may cause an effect in the cleavage site of Spike^Omicron^, the decreasing flexibility in relation to Spike^WT^, may lead to higher viral infectivity due to better exposition of the furin cleavage site in this variant. As D614G mutation is present in all VOCs, the decreasing flexibility may explain the increased efficiency of Spike cleavage by furin observed for many VOCs [27–29]. Thus, by using molecular dynamics simulations, we have detected various structural changes in the Spike^Omicron^ ectodomain that help explain features such as the increase in viral infectivity of this variant, while revealing aspects that may be relevant for immune system evasion. Further simulations exploring details of the interaction of this variant with hACE2 and both computational prediction and experimental scanning of epitopes will help clarify the impact of such structural changes in the interactions of the virus and its host.

## Acknowledgment

The authors acknowledge the National Council for Scientific and Technological Development (CNPq), the Coordination for the Improvement of Higher Education Personnel (CAPES, grant 88887.374931/2019-00, Coordenação de Aperfeiçoamento de Pessoal de Nível Superior - Finance Code 01), and the São Paulo Research Foundation (FAPESP, grants 2019/00195-2, 2020/04680-0 and 2016/09047-8), Rede Virus MCTI (grant FINEP 0459/20), Brazil, for financial support.

